# A tail of two sides: Artificially doubled false positive rates in neuroimaging due to the sidedness choice with *t*-tests

**DOI:** 10.1101/328567

**Authors:** Gang Chen, Robert W. Cox, Daniel R. Glen, Justin K. Rajendra, Richard C. Reynolds, Paul A. Taylor

## Abstract

One-sided *t*-tests are widely used in neuroimaging data analysis. While such a test may be applicable when investigating specific regions and prior information about directionality is present, we argue here that it is often mis-applied, with severe consequences for false positive rate (FPR) control. Conceptually, a pair of one-sided *t*-tests conducted in tandem (e.g., to test separately for both positive and negative effects), effectively amounts to a two-sided *t*-test. However, replacing the two-sided test with a pair of one-sided tests without multiple comparisons correction essentially doubles the intended FPR of statements made about the same study; that is, the actual family-wise error (FWE) of results at the whole brain level would be 10% instead of the 5% intended by the researcher. Therefore, we strongly recommend that, unless otherwise explicitly justified, two-sided *t*-tests be applied instead of two simultaneous one-sided *t*-tests.

## INTRODUCTION

The *t*-statistic is a dimensionless ratio that indicates the effect estimate relative to the corresponding standard error of the effect estimate. As a result, it provides the assessment in making inference for an effect in a model, with common examples in neuroimaging being one-sample tests, two-sample tests, paired tests, general linear models (GLMs), linear mixed effects (LME) models, etc. Under the null hypothesis significance testing (NHST) framework, one first formulates a null hypothesis for an effect of interest μ (e.g., the intercept or slope in a model) such as “*H*_o_: μ = 0,” and then there would typically be three possible alternative hypotheses: one two-sided,^1^ “*H*_a_: μ ≠ 0,” and two one-sided, “*H*_a_: μ > 0” and “*H*_a_: μ < 0.” Once the alternative hypothesis is chosen, the statistical formulation is often clearly defined before the real data analysis occurs, and therefore the testing formula itself does not involve the issue of sidedness. Indeed, the null and alternative hypotheses should provide the initial impetus for the experiment itself and directly determine its design as well as any modeling or testing.

While the above description about such a common testing strategy may be familiar to researchers and data analysts, we find that unfortunately there is persistent confusion and pervasive mishandling in the neuroimaging field about the appropriate way to choose between one- and two-sided testing. Importantly, such mis-testing often leads to inflated false positive rates (FPR) and familywise error (FWE). For example, one might reasonably think to apply a pair of one-sided tests in order to separate the directionality of estimated effects. However, as we discuss below, this actually doubles the expected FPR at the voxel level as well as the familywise error (FWE) rates at the whole brain level, and there are other direct and unproblematic ways of separating directionality.

Here, we first aim to clarify between cases where one-sided or two-sided testing is appropriate. The general rule can actually be stated succinctly: *If the investigator can justify focusing on one side (direction) based on prior knowledge, then they should explicitly state their reasoning and decision,*^*2*^ *and perform their individual one-sided test; otherwise, two-sided testing should be conducted.* This can be applied straightforwardly to voxelwise, ROI-based or other study approaches, as described below.

We then demonstrate the necessity of following the above rule and the statistical problems with using a pair of one-sided tests instead of a single two-sided test. Unfortunately, sidedness malpractice is quite common in the neuroimaging field, even when addressing a two-sided hypothesis and alarmingly *without* performing appropriate multiple comparisons corrections. We describe why such a practice is both unnecessary and statistically misleading, and how it introduces two forms of FPR inflation in neuroimaging studies. We make recommendations for clear analysis procedures and reporting, and we hope that these proper statistical practices are adopted in the field.

## BACKGROUND THEORY AND FOREGROUND PRACTICE

In a scientific study, a standard form of question to ask under NHST is, “For some given sample data, what can one say about the population effect being different than zero?” When the sign of the effect is unknown in advance, this would be addressed by using a *t-*test as follows: set up the null hypothesis *H*_o_: μ = 0 and an alternative hypothesis *H*_*a*_: μ ≠ 0; calculate a statistic for the sample; and test it against a nominal significance threshold level, say *α*. Importantly, by looking for *any* difference, the NHST is performed by checking whether the effect is “statistically different” from zero on either the positive *or* the negative side at a preset total significance level *α* (i.e., the sum of the two tails): a single hypothesis is investigated by checking both tails of a distribution. Thus, the phrasing of the question tautologically dictates that two-sided testing is appropriate. For symmetric distributions, such as a Gaussian and Student’s *t*, the FPR is evenly split between the two tails (i.e., the area under each tail is *α*/2).

If the sign of the effect is known or strongly suspected *a priori*, then a one-sided test can certainly be justified. For example, if previous studies indicate that the effect is likely positive (or likely negative), a testing platform can be constructed as *H*_o_: μ = 0 vs *H*_*a*_: μ > 0 (or vs *H*_*a*_: μ < 0), where the FPR is controlled through the designated directionality; that is, the area under the preselected tail is *α*. In this case, a one-sided test would be reasonable, and the researcher must weigh the benefit of having the full sensitivity with an FPR of *α* in a single direction versus conducting a two-sided test and having lower sensitivity with an FPR of *α*/2 in both directions but perhaps not missing a possible detection in the other direction.^3^

However, an all-too common malpractice in neuroimaging is to adopt a hybrid testing strategy of the above: to conduct a pair of one-sided *t*-tests simultaneously (*H*_*a*1_: μ > 0 and *H*_*a*2_: μ < 0), and then to report/display those results separately. This dual testing is unnecessary and inefficient in theory, and unfortunately it tends to be implemented incorrectly and spuriously in practice.

First of all, substituting a pair of one-sided tests cannot be justified simply because the investigator would like to reveal the directionality of an effect.^4^ Two-sided testing achieves the same goal through the sign of either the effect estimate or the *t*-statistic value. Therefore, having two separate tests does not add any information to the results (while it *would* necessitate multiple testing correction). The purpose of NHST is to decide to whether to reject *H*_*o*_ or not; the determination of an effect’s directionality comes from the posterior information that is present in the result, not from the prior choice of sidedness in test formulation.

On the other hand, substituting an *F*-test, which is conceptually equivalent to a two-sided test and implemented in its place by some softwares, also has difficulties. The *F*-test with one numerator degree of freedom is equal to the *t*-value squared of the two-sided test (with the same denominator degrees of freedom). However, the typical two-sided *t*-test actually provides posterior information about the directionality of the effect estimate, while the corresponding *F*-test hides that information. A separate calculation outside of the *F*-test would still be required for that information (but, as discussed above, using a pair of one-sided tests to do this would lead to FPR inflation, so other means would be required). In summary, a two-sided test is more informative, much simpler and avoids FPR inflation pitfalls.

How does the above extend to the neuroimaging case of whole brain or region of interest (ROI) studies? Consider two common scenarios of neuroimaging group analysis:

A. A researcher performs whole brain analysis with a massively univariate approach, where the same model is applied to each voxel. All hypotheses are then shared across the brain, and it is difficult to imagine a scenario where this researcher would test all voxels simultaneously for just one directional change. It is much more likely they would look for differences of either side, making this a case for two-sided testing in essentially all circumstances.
B. When performing region-wise or more localized analyses, it is conceivable that a researcher may have background information to expect changes of a certain direction in some set of regions *R*_1_ and of the other direction in set *R*_2_. In this scenario it might make sense to apply individual one-sided tests within the regions of *R*_1_ and *R*_2_, as long as the two sets are independent.^5^

In both cases, the same basic rule applies when performing the same test across the whole brain: *if the* a priori *information does not lead a researcher to a hypothesis of the same one-sided change (such as H*_*a*_: *μ > 0) across the domain examined, then two-sided testing must be uniformly performed.* If the researcher can satisfactorily justify examining one direction of change for one or more regions, then they can reasonably apply a single one-sided test in those locations. (Additionally, as a separate consideration, if the researcher is analyzing several voxels or regions simultaneously, they would also be expected to perform multiple testing correction for their FPR across those voxels or regions, discussed further below.)

## FPR CONTROLLABILITY PROBLEMS WITH PAIRS OF ONE-SIDED TESTS

What are the consequences of performing a pair of one-sided tests in place of a two-sided test? While the combination of “*H*_*a*_: *μ* > 0 or *μ* < 0” is logically equivalent to ”*H*_*a*_: *μ* ≠ 0,” in terms of a statistical framework, splitting the single two-sided test into two one-sided tests changes FPR controllability, as we now explore.

The FPR of hypothesis rejection is controlled by the total designated area in the statistical tail(s), *α.* Because of the symmetry of the *t*-distribution, performing two-sided testing with an overall significance level of *α* has exactly equivalent results to performing a pair of one-sided tests at the significance level of *α*/2. In other words: *performing a pair of one-sided tests at the significance level of α is equivalent to performing two-sided testing at the significance level of 2α, and therefore the expected FPR is actually twice the stated rate.* This intrinsic relation, and the fundamental factor-of-two difference between a two-sided test at *α* and a pair of one-sided tests each at *α*, is shown in Fig. 1.

**Figure 1.**
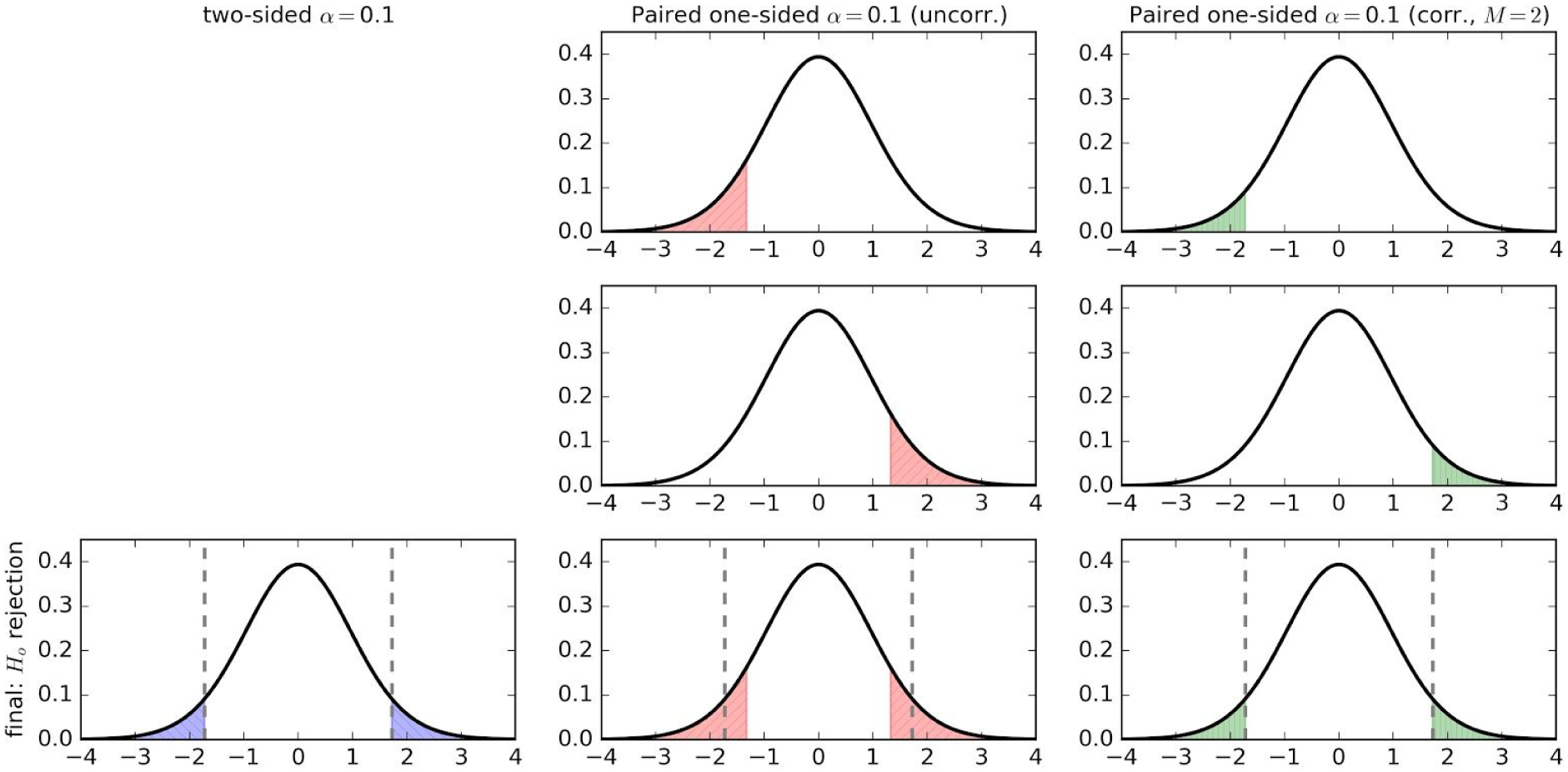
Consider testing a null hypothesis *H*_*o*_: μ = 0 versus an alternative hypothesis H_a_: μ ≠ 0 for the case of a Student *t*_20_*-*distribution. Comparisons of null hypothesis rejection (shaded regions, which represent FPR) for each test at the same reported significance level (*α* = 0.1, for visibility of the areas) are shown in the bottom row: (column 1) two-sided testing at *α* = 0.1, threshold at ±1.72; (column 2) unadjusted pairs of one-sided tests each at *α* = 0.1 (with the combined FPR shown in the bottom row), threshold at ±1.32; and (column 3) two one-sided tests each at *α*=0.05 (with the combined FPR shown in the bottom row), threshold at ±1.72. Vertical dashed lines show the boundaries of the appropriate two-sided testing, for comparison. The unadjusted pair of one-sided tests have a much larger area (by a factor of 2) of null hypothesis rejection, necessarily translating into doubled FPR.

One may also view this as a case of Bonferroni correction for *M* = 2 multiple tests when conducting the pair of one-sided tests, which are necessarily orthogonal (making the correction exact). The problem arises in neuroimaging that in many cases of paired one-sided testing the sidedness corrections are *not* performed, so that one would necessarily expect the ensuing results to have higher FPR (by a factor of 2; see again Fig. 1) than the targeted value.

This sidedness issue is not a matter of interpretation about effect directionality but simply a black-and-white, quantitative fact about FPR controllability. One way to observe this is to compare plots of relative areas of null hypothesis rejection for a given threshold, as demonstrated in Fig. 1, where the significance level *α*=0.1 is applied to an example of Student’s *t*_20_-distribution. In the first column, the area (= FPR) showing where two-sided testing would reject the null hypothesis is shown in blue. The second columns shows the FPR of each of the pair of one-sided tests, as well as their combined FPR at the bottom in red; the total FPR of rejecting the null hypothesis is much larger than in the two-sided case (by a factor of two, as shown above). The third column shows how adjusting appropriately for *M* = *2* comparisons restores the pair of one-sided tests to agreement with the two-sided test.

One might also compare the threshold of a statistic associated with a given significance level *α*, since neuroimaging researchers often translate a voxel-wise significance level for the purpose of thresholding a statistic map as the first step toward overall FPR in the whole brain. For a given significance level *α* (say, 0.001) and number of degrees of freedom (say, 20), one can compare the associated *t*-statistic threshold under the various scenarios. In the two-sided case, one finds *t*_20_ *≈* 3.90, and in a one-sided case, *t*_20_ *≈* 3.55, a lower threshold as expected from viewing Fig. 1. When two one-sided tests are performed, *and* the appropriate adjustment is applied, then the adjusted voxel-wise significance level is 0.001/2 = 0.0005, and the corresponding *t*_20_ *≈* 3.90 is in agreement with the original two-sided test.

## APPLICATION TO FMRI DATA: DOUBLING OF INTENDED FWE

The principle stated above, that the practice of simultaneously performing a pair of one-sided tests doubles the intended FPR, applies in any context. Of particular interest here is its application to neuroimaging studies, where multiple testing correction is applied at the whole brain level (for example, to maintain an overall FWE of 5%). Regardless of whatever correction procedure one adopts -- Monte Carlo simulations, random field theory, threshold-free cluster enhancement (TFCE), etc. -- a pair of one-sided tests essentially leads to an effective overall FWE of 10% after the correction, not to the intended level of 5%.

We now demonstrate how the above theoretical features affect an FMRI study in practice, by performing a group-level analysis on a set of real data.^6^ We compare the results of a pair of one-sided tests, which would require multiple testing correction at both voxel and whole brain levels, with a two-sided test. The task fMRI data used here comes from OpenfMRI (Poldrack et al., 2013), using a group of 16 subjects from ds000001 (release 2.0.4). It was processed using AFNI (Cox, 1996) following the “NIMH” set of steps described in Taylor et al. (2018), with single-subject processing including blocks for motion correction, alignment to standard space, etc. (also note the comments about processing steps described therein; for the present purposes of showing statistical effects, the data are reasonable). The group analysis was performed using 3dMEMA for voxelwise mixed-effects multilevel analysis (Chen et al., 2012) with both the effect estimate and the corresponding *t*-statistic at the individual subject level as inputs.

Here, using the average spatial smoothness parameters of the group with the mixed-model autocorrelation function (ACF), 3dClustSim is executed to create cluster tables, so that one obtains a cluster size threshold *C* = *C(p, α)* for a given voxelwise threshold *p* and nominal (whole brain) FWE *α*. Tables are automatically created for three possible definitions of a voxel’s neighborhood number (NN): NN = 1, facewise neighbors (6 total); NN = 2, face+edgewise neighbors (18 total); and NN = 3, face+edge+nodewise neighbors (26 total).^7^ Additionally, tables are created for three separate forms of cluster sidedness:

- one-sided: minimum *C* after voxels are thresholded > *t*(*p*) in order to achieve FWE = *α* for positive effects; the same applies to thresholding < -*t*(*p*) for negative effects.
- two-sided: minimum *C* after voxels are thresholded > *t*(*p/*2) or < -*t*(*p/*2), in order to achieve FWE = *α*; here, a single cluster could be comprised of both positive and negative effects.
- bi-sided (another form of two-sided): minimum *C* after voxels are thresholded at *t*(*p/*2) or < *-t*(*p/*2), in order to achieve FWE = *α*; here, each single cluster is of entirely positive or entirely negative effects.

That is, bi-sided clustering is the same as two-sided, with the additional restriction of separating voxels with different signs of effect. In general, it seems reasonable that bi-sided clustering behavior would be the desired form for most neuroimaging questions (though we also note that, in practice, the smoothness characteristics of neuroimaging data lead to near equivalence of bi- and two-sided clusterization).

An example cluster table for these data is shown in Fig. 2 (one-sided test and NN = 1). After specifying a voxelwise threshold *p* and overall FWE *α*, one finds the cluster size threshold *C* (e.g., using the ceiling of the table value, which may be non-integer due to the interpolation of simulation results, so 50 here). One would apply the same (one-sided) voxelwise threshold to the actual statistic dataset, and then keep only (NN = 1) clusters with ≥ *C* voxels. In this way the remaining voxel clusters should represent a dataset with overall FWE of *α*, with the same steps applying regardless of test sidedness or voxel neighbors.

**Figure 2.**
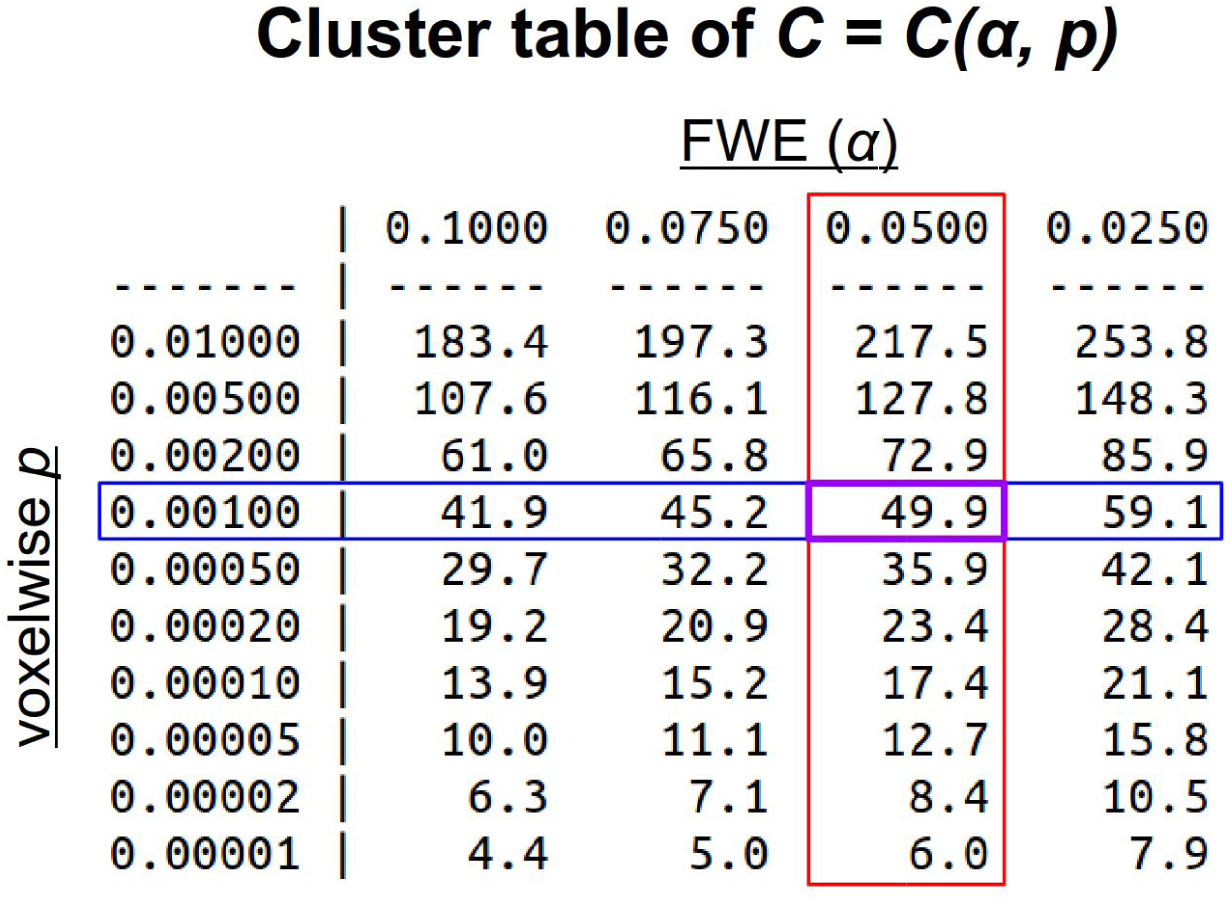
An example cluster table generated by 3dClustSim for a given NN and sidedness of testing. The user specifies a desired voxelwise threshold *p*-value (here, *p* = 0.001, in blue) and desired nominal FWE rate (here, *α* = 0.05, in red), and then finds the cluster threshold *C*, which is in units of number of voxels, associated with that value (purple). The threshold value can be recorded as a decimal due to interpolation of simulation results; here, the example cluster size threshold for chosen parameters would be *C* = 50 voxels.

We now examine clustering results iexamine clustering results innexamine clustering results in the analyzed data set, comparing bi-sided and one-sided results using NN = 1 (results are equivalent across NN values). Here, we investigate clustering results for common values of voxelwise threshold *p*=0.001 and nominal FWE *α* = 0.05.

First, we compare the cluster tables and investigate the following question: if performing clustering for a pair of one-sided tests with *p* = 0.001 and *α* = 0.05, what is the equivalent FWE for a single bi-sided^8^ test? The translation of results is shown in Fig. 3. The one-sided tests’ *p*_1_= 0.001 translates to a bi-sided *p*_2_ = 0.002 (see, e.g., Fig. 1), so we find the bi-sided cluster value that approximately equals that of the one-sided result, *C*_2_ ≈ *C*_1_. Here, that is 49.6, which is in the *α* = 0.10 column. Thus, as predicted, using a pair of one-sided tests instead of a bi-sided test doubles the FWE for the reported whole brain results. Reporting results from a pair of one-sided tests with any *p = X* and *α* = *Y* is essentially equivalent to those of a bi-sided test with *p = 2X* and *α* = 2*Y*.

**Figure 3.**
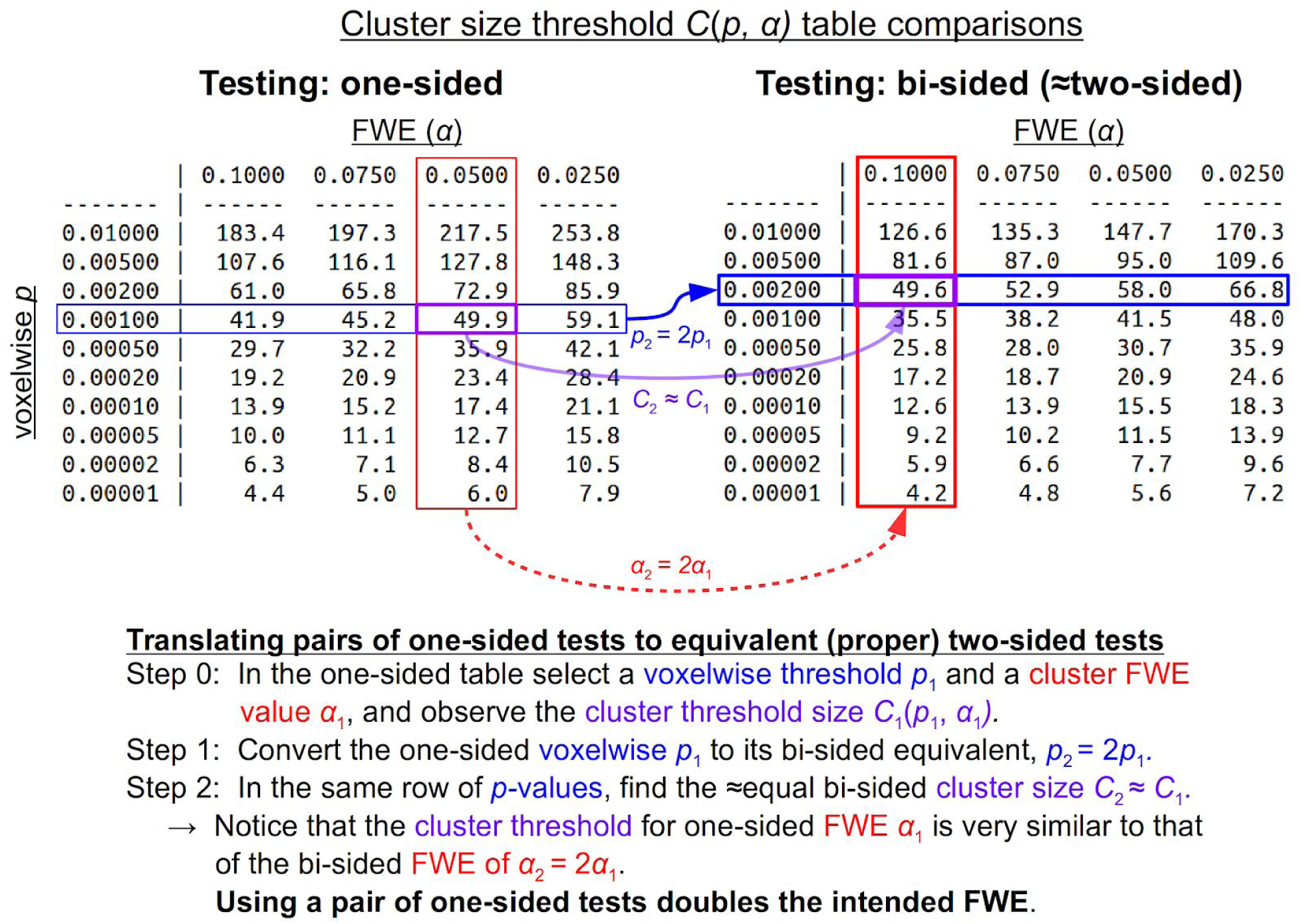
Schematic demonstrating how cluster-level results for pairs of one-sided tests are interpretable in terms of proper two-sided testing (from the ds000001 dataset examined here; see Fig. 4 for related brain images, and Table 1 for summarizing cluster information). When a researcher chooses a particular voxelwise threshold (here, *p* = 0.001) and FWE value (here, *α* = 0.05), they obtain a corresponding cluster size *C*, which serves as a cluster-level threshold. As shown here, because of the multiple testing involved in having a pair of one-sided tests, the reported FWE *α* is actually much larger than reported (essentially by a factor of 2). Note that this equivalency to a double FWE occurs throughout these ranges of *p*- and α-values, and applies generically in the same manner to other software implementations.

Fig. 4 shows the effects of this phenomenon in the actual data. Row A displays the results of bi-sided clustering, *p* = 0.001 and *α* = 0.05.^9^ There were 19 clusters found, containing a total of 5337 voxels (see Table 1). Notice as well that the directionality of each cluster is clear from the effect estimate. Row B of Fig. 4 shows the results of a pair of one-sided tests with the same voxelwise and FWE parameters; there were 22 clusters found over a total of 8148 voxels (differences highlighted in the figure) due to the doubled FWE. In row C, the voxelwise threshold has been adjusted for the pair of one-sided tests, but the FWE is still effectively doubled and more clusters are still observed. When both the voxelwise threshold and the FWE values are adjusted for the pair of one-sided tests, row D shows that the results are equivalent to those of the correct bi-sided test.

**Table 1.**
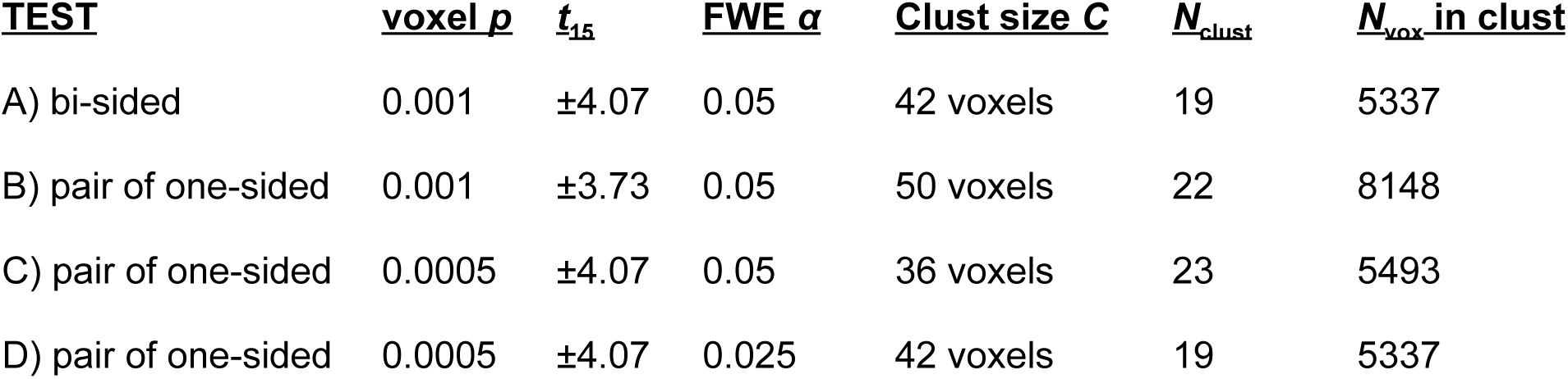
Tabulated results of the clustering process from the ds000001 dataset analyzed here (see Fig. 3 for cluster table information and Fig. 4 for results in the brain). For each test, a given voxelwise threshold *p* (which can be converted to a *t*-statistic for a given number of degrees of freedom) and cluster-level FWE *α* to obtain a cluster size threshold *C* from the results of 3dClustSim. The voxelwise *p* and cluster size *C* thresholds are then applied to the data of interest, producing final maps of clusters of interest that pass the threshold. Here, we report the number of clusters *N*_clust_ and the total number of voxels summed across these clusters *N*_vox_for each of the testing scenarios. Only the one-sided tests with adjustments applied to both *p* and *α* (D) produce results matching the bi-sided case. The other one-sided tests cases (B and C) have uniformly larger numbers of cluster and voxels in clusters, particularly B, which corresponds to many cases in existing literature.

| **<u>TEST</u>** | **<u>voxel $p$</u>** | **<u>$t_{15}$</u>** | **<u>FWE $\alpha$</u>** | **<u>Clust size $C$</u>** | **<u>$N_{\text{clust}}$</u>** | **<u>$N_{\text{vox}}$ in clust</u>** |
| --- | --- | --- | --- | --- | --- | --- |
| A) bi-sided | 0.001 | $\pm 4.07$ | 0.05 | 42 voxels | 19 | 5337 |
| B) pair of one-sided | 0.001 | $\pm 3.73$ | 0.05 | 50 voxels | 22 | 8148 |
| C) pair of one-sided | 0.0005 | $\pm 4.07$ | 0.05 | 36 voxels | 23 | 5493 |
| D) pair of one-sided | 0.0005 | $\pm 4.07$ | 0.025 | 42 voxels | 19 | 5337 |

**Figure 4.**
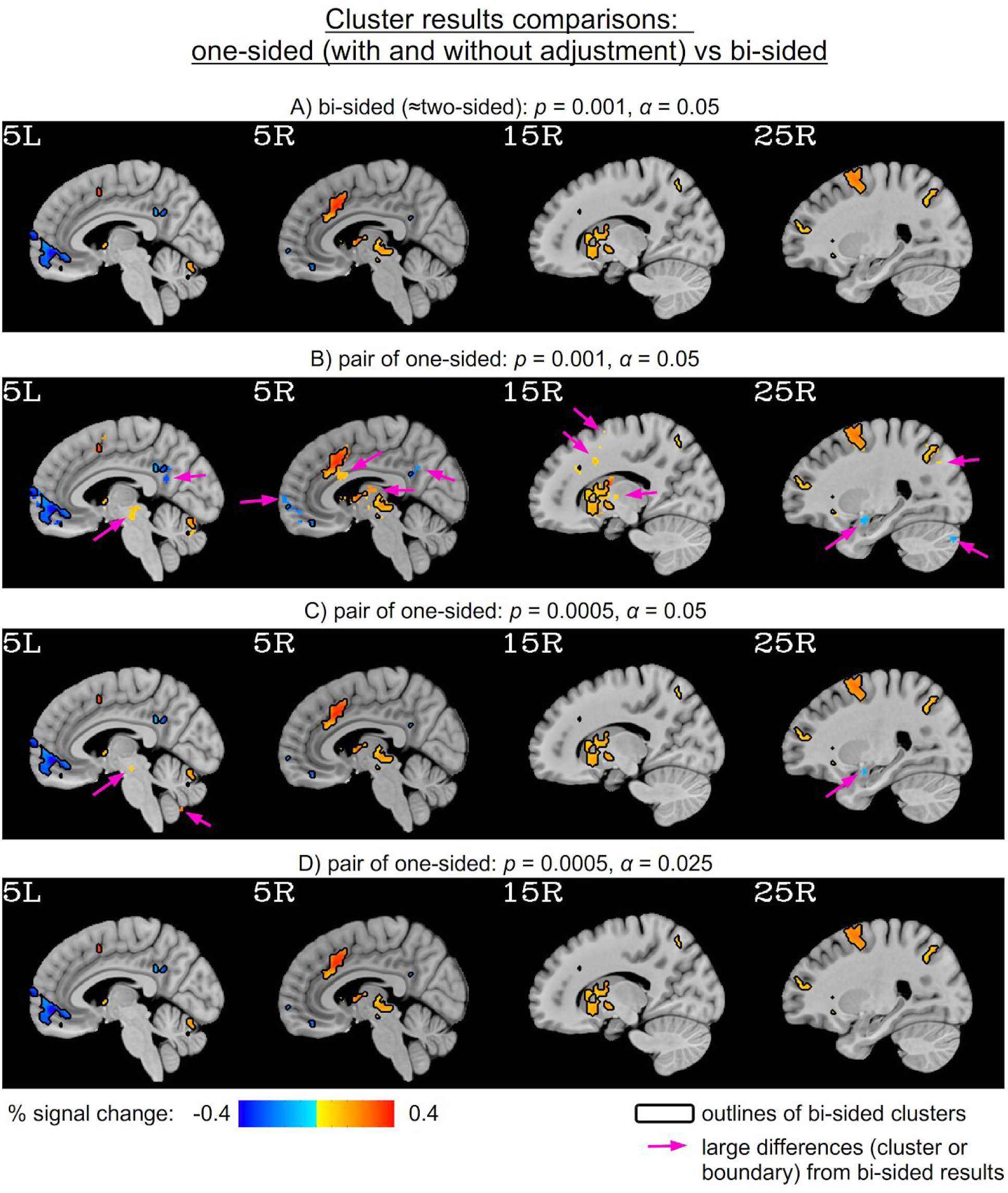
Group-level results from ds000001 comparing results with correct bi-sided testing (top row) with pairs of one-sided testing (cluster counts summarized in Table 1). In each case, the bi-sided results are shown with black outlines, for reference, and the pink arrows highlight large differences (full clusters or cluster boundary variation) from the bi-sided results. Displayed overlay values show the effect estimates of the data in units of BOLD %-signal change; from these effects, directionality is apparent (as well as from the sign of the associated statistic values). The second row shows one-sided testing results using the same reported voxelwise threshold p and cluster FWE *α* used for the bisided; the threshold-passing regions show many differences from the bi-sided results, as expected. In the third row the voxelwise parameter p has been adjusted to the same value as bisided; while the differences are smaller than in the unadjusted case, there are still more clusters reported. In bottom row, both the voxelwise threshold and the FWE have been adjusted, and the cluster results essentially match those of the bi-sided tests. Each of these results are expected from the theoretical explanations presented in the text.

## DISCUSSION AND CONCLUSION

In summary, we argue that the neuroimaging field should adopt a clear and consistent approach to dealing with the sidedness of *t*-tests. We have focused on FMRI here, but the same issue applies to any area of study, such as DTI, PET, EEG, etc. Regardless of modeling perspective (conventional, Bayesian, etc.), spatial domain (voxelwise, ROI-based, cross-regional, etc.), or grouping (one-sample, two-sample, etc.), researchers should be careful about the statistical testing that they choose, ensure that it matches their stated hypotheses, and be precise about the way that they report it. The chosen sidedness of testing should always be made clear, because just reporting the *α* and/or *p*-value is not enough information. The setup of the hypothesis should be directly justified, and the sidedness of the test should be consistent with it so that the purported controllability (false positive rate or posterior probability) matches the intended goal. Arguments for utilizing a one-sided test can be made if prior knowledge of directionality is available for all regions together. Otherwise, two-sided testing is more generally appropriate.

Within the field of neuroimaging, there have been questions raised about maintaining overall FWE control in studies, particularly related to user-chosen parameters like the voxelwise one-sided *p* (Eklund et al., 2016).^10^ Specifically, higher voxelwise one-sided *p*-values (∼0.01) were associated with worse FWE control (with some additional dependence on type of task, number of samples and other parameters) in those tests (Eklund et al., 2016). Those findings created a strong discussion in the field, and even led many reviewers and some journal editors (Roisier et al., 2016) to reject papers that implement thresholding at voxelwise significance level of 0.01. It is not clear that those simulation results could be extended to the situation of two-sided testing, since Eklund et al. (2016) only used one-sided tests in their simulations. However, the issue raised in the present study is even more fundamental to FWE control than those previous studies, and we hope that editors and reviewers alike will join us in asking for statistical correctness and clarity in published studies.

As recently reported in Table 1 of Pauli et al. (2016), the default test type in AFNI is two-sided (though, as discussed above, the closely related “bi-sided” is typically recommended), and one-sided testing is easily available, when appropriate. However, one-sided testing was reported as the default for both FSL and SPM, with two-sided testing listed as “NA“. Without a two-sided test directly available, one would have to finesse the statistical testing by using an *F*-test and/or pairs of one-sided tests with adjusted values (correcting both voxelwise and cluster-wise threshold to *p/*2 and *α*/2, respectively).

Within FMRI research, most people do not report the sidedness of their testing; we can therefore only guess the sidedness of their testing, but it would likely be the default of their software. Searching in Google Scholar, there were 670,000 results for pages with FMRI and no inclusion of forms of one-sided/tailed, two-sided/tailed or *F*-test.^11^ There were 7,560 with FMRI and explicit one-sided/tailed testing^12^ (though, we did not check how many were whole brain vs how many were possibly ROI-based with one-sided hypotheses), and only 18,700 results for pages with FMRI and explicit two-sided/tailed testing.^13^ In summary, a large fraction of studies, likely even the majority, appear to be using one-sided testing.^14^

We would recommend that journals require explicit information about sidedness when reporting results. Instead of reporting that clusters were generated using, e.g., “a voxelwise threshold of *p =* 0.001 and FWE = 5%“, authors should state, “a voxelwise threshold of two-sided *p =* 0.001, with 49 degrees of freedom and FWE = 5%.” Furthermore, *if* one-sided testing is applied, then it should be explicitly and carefully justified (e.g., by prior research, previous test studies, etc.) and/or clearly localized. Finally, pairs of one-sided tests over the same domain should simply not be performed -- there does not appear to be any valid reason for such, and such testing can easily be abused and misused.

## ACKNOWLEDGMENTS

The research and writing of the paper were supported by the NIMH and NINDS Intramural Research Programs (ZICMH002888) of the NIH/HHS, USA. This work utilized the computational resources of the NIH HPC Biowulf cluster. (http://hpc.nih.gov).

## Footnotes

1 One- and two-sided hypotheses are sometimes referred to as one- and two-tailed hypotheses, respectively. “Tail” under NHST indicates the far end area (or FPR) under the probability density function, but we prefer “sided” for its connotation of directionality.

2 Reporting and justifying sidedness, together with the degrees of freedom for *t*-test, is largely lacking in neuroimaging literature.

3 We note that this reasoning and all arguments below basically applies to *all t-*tests: comparing differences of groups, testing differences from a constant *K* via *H*_o_: μ = *K* and *H*_*a*_: μ ≠ *K*, etc.

4 On a note of full disclosure, one of these authors (PT) has previously used pairs of one-sided tests for just such a purpose, but now hopes that others will be spared the blushes from this fallacy.

5 For instance, a decision at the regional level can be adopted in an ROI-based analysis approach through Bayesian multilevel modeling (Chen et al., 2018).

6 The group-level processing script (running 3dMEMA, 3dClustSim, 3dClusterize, etc.) for ds000001 used here is available at: https://afni.nimh.nih.gov/pub/dist/doc/htmldoc/codex/fmri_brief.html

7 Results are equivalent across NN values, as long as the same NN is used across a single analysis

8 Again, here and in most neuroimaging contexts, bi-sided and two-sided results would be essentially identical. In the scenario that they would not be, typically the interpretation of bi-sided clusters seems more logical for neuroimaging interest.

9 Both the cluster tables and the cluster maps were identical for the two-sided and bi-sided results here, which is common in FMRI data with standard smoothing applied, as noted above.

10 The cluster-level performance of overall FWE at the cluster level, as assessed through simulations in Eklund et al. (2016), only addresses the FWE controllability for each individual one-sided test; in other words, the inflation factor of two in overall FWE remains whenever a pair of one-sided tests is adopted.

11 Google scholar search string: *intext:fmri -“one-tailed” -“one-tail” -“one-sided” -“1-sided” -“two-tailed“ -“two-tail” -“one-sided” -“2-sided” -“F-test“*.

12 Google scholar search string: *intext:fmri -“two-tailed” -“two-tail” -“one-sided” -“2-sided” -“F-test” intext:“one-tailed” OR intext:“one-tail” OR intext:“one-sided” OR intext:“1-sided“.*

13 Google scholar search string: *intext:fmri -“one-tailed” -“one-tail” -“one-sided” -“1-sided” intext:“two-tailed” OR intext:“two-tail” OR intext:“one-sided” OR intext:“2-sided” OR intext:“F-test“.*

14 The commonality of this situation is highlighted by an anecdote: during a recent manuscript review process of ours, the statement, “testing both tails in tandem… is widely used” (Chen et al., 2018) evoked an unexpectedly dismissive denial from a reviewer: “Really? I disagree. Some reference or quantification required.”

